# Unappreciated Subcontinental Admixture in Europeans and European Americans: Implications for Genetic Epidemiology Studies

**DOI:** 10.1101/2022.11.28.518227

**Authors:** Mateus H. Gouveia, Amy R. Bentley, Eduardo Tarazona-Santos, Carlos D. Bustamante, Adebowale A. Adeyemo, Charles N. Rotimi, Daniel Shriner

## Abstract

European-ancestry populations are recognized as stratified but not as admixed, implying that residual confounding by locus-specific ancestry can affect studies of association, polygenic adaptation, and polygenic risk scores. We integrated individual-level genome-wide data from ^~^19,000 European-ancestry individuals across 79 European populations and five European American cohorts. We generated a new reference panel that captures ancestral diversity missed by both the 1000 Genomes and Human Genome Diversity Projects. Both Europeans and European-Americans are admixed at subcontinental level, with admixture dates differing among subgroups of European Americans. After adjustment for both genome-wide and locus-specific ancestry, associations between a highly differentiated variant in *LCT* (rs4988235) and height or LDL-cholesterol were confirmed to be false positives whereas the association between *LCT* and body mass index was genuine. We provide formal evidence of subcontinental admixture in individuals with European ancestry, which, if not properly accounted for, can produce spurious results in genetic epidemiology studies.

## INTRODUCTION

Human genetic studies have primarily considered admixed populations to have resulted from interbreeding between two or more continentally separated populations^1–3^. However, continental ancestry is not necessarily a single homogenous component of genetic diversity, but rather can be a composite of diverse subcontinental ancestries^4,5^. In some instances, differentiation between intra-continental populations is on par with or higher than differentiation between inter-continental populations^1,6^. Also, there are examples from pharmacogenetics of variants that are differentiated at the intra-continental level, such as in the case of abacavir hypersensitivity syndrome, for which the causal allele (HLA-B*5701) has a prevalence of 13.6% among Maasai in Kenya but a prevalence of ^~^0% among Yoruba in Nigeria^7^.

Despite genetic studies highlighting a clear pattern of North-to-South genetic variation in Europe^8–10^ and strong evidence of admixture within Europe by ancient DNA analysis^11,12^, European-ancestry populations are generally treated in association models as stratified but not as admixed at the subcontinental level. As a result, genetic epidemiology studies of Europeans or European Americans usually control for potential confounding effects of population stratification using genome-wide ancestry estimated by principal components analysis^13^, but do not control for locus-specific ancestry, which is inherent to admixed populations^14^. Potential consequences are that detection of causal genetic variation is hampered and estimation of effect sizes can be biased, leading to further negative consequences such as misestimation of polygenic adaptation^15^ and poor predictive performance of polygenic risk scores^16^.

Recently developed approaches have enabled the use of genome-wide data (either array-based genotype or whole genome sequence data) to assess admixture at two levels: genome-wide ancestry (also known as global ancestry)^13,17,18^, which is the individual’s ancestry averaged across the entire genome, and locus-specific ancestry (also known as local ancestry)^19^, which allows for inference of an individual’s ancestry at each locus. The power, resolution, and specificity of disease or trait mapping studies can be improved by leveraging both genome-wide and locus-specific ancestries^3,20,21^. To assess both genome-wide and locus-specific ancestries in admixed individuals, present-day populations are used as proxies for ancestral populations that serve as references for ancestry estimation. Considering that ^~^96% of participants in genome-wide association studies (GWAS) have European ancestry^22^, a comprehensive analysis is needed to evaluate the adequacy of European reference panels for ancestry analysis using European-ancestry individuals.

The prevalence of lactase persistence varies widely across Europe and the most strongly associated variant rs4988235 in the lactase gene (*LCT*) has been reported to be under positive selection and associated with height, body mass index (BMI), and low-density lipoprotein (LDL)^23–26^. The SNP rs4988235 is one of the most highly differentiated variants in Europe^27^, with derived allele (A) frequencies ranging from 93.1% in Swedes to 2.9% in Sardinians^28^. Importantly, rs4988235 and height are well-known to covary following a north-to-south axis^29^, and the association between rs4988235 and height has been suggested to be spurious based on attenuation following adjustment for genome-wide ancestry^25^. Nonetheless, there are no association studies in European-ancestry populations that control for confounding at both the genome-wide and locus-specific ancestry levels to test the validity of the association between rs4988235 and reported associated traits.

To test for the existence of subcontinental ancestries within Europe, we integrated genome-wide data from 1,216 individuals across 79 European populations. Then, to examine population structure and admixture, we integrated genome-wide data from 17,669 European Americans from five genetic epidemiology cohorts in the US. Finally, to illustrate the potential implications of confounding by subcontinental ancestry and admixture, we interrogated the association between rs4988235 and height, LDL-cholesterol, and BMI.

We found that the 1000 Genomes and Human Genome Diversity Projects provided incomplete coverage of European ancestries, so we generated a new reference panel to capture additional European ancestral diversity. Our admixture analyses yielded formal evidence that European-ancestry individuals are admixed at the subcontinental level, with admixture dates differing among European Americans. After adjustment for both genome-wide and locus-specific ancestry, previously reported associations between rs4988235 and height or LDL were no longer statistically significant, strongly supporting that they are false positives due to uncorrected stratification. We observed systematically better fits when models were adjusted for principal components (PCs) derived from projection of European Americans onto our new reference panel, rather than for PCs derived from study-specific unsupervised analysis. Altogether, this study indicates that full adjustment for subcontinental European admixture (at both genome-wide and locus-specific levels) should become best practice in genetic association studies using European-ancestry individuals, including the UK Biobank^30^ in Europe and the All of Us Research Program^31^ and the Million Veteran Program^32^ in the United States.

## RESULTS

### Reference panels of European diversity

We generated a new reference panel capturing genetic diversity from 79 European populations from five population genetics studies: the 1000 Genomes Project^33^, the Human Genome Diversity Project (HGDP)^34^, the Human Origins dataset^35^, a study of the Caucasus Mountains^36^, and a study of the Jewish Diaspora^37^ (Fig. 1A and Table S1). After quality control to reduce batch effects, our European panel included 1,216 unrelated individuals and 104,414 genotyped SNPs. Principal component analysis (PCA)^13^ showed that North Europeans (*e.g*., Finnish, Lithuanian, and Estonian) *vs* Southeast Europeans (*e.g*., Armenian, Georgian Jew, and Georgian Megrel) represented the extremes along the first principal component (Fig. 1B). Along the second principal component, Southwest Europeans (*e.g*., Sardinian, Basque, and Spanish) *vs* Southeast Europeans (*e.g*., South Caucasus) represented the extremes. Subsequent principal components separated population-specific genetic variability (Fig. S1). To compare our panel with commonly used European reference panels from the Human Genome Diversity Project (HGDP)^34^ and the 1000 Genomes Project^33,34^, we calculated convex hull areas^38^ defined by the first two principal components (Fig. 1B and 1C). Compared to our panel, the 1000 Genomes Project and the HGDP covered 26.8% and 61.3% of European diversity, respectively, while the combination of the 1000 Genomes Project and the HGDP covered 77.3% (Fig. 1C). These results indicate that the 1000 Genomes Project and the HGDP, separately and combined, provide incomplete coverage of European genetic diversity.

**Fig. 1.**
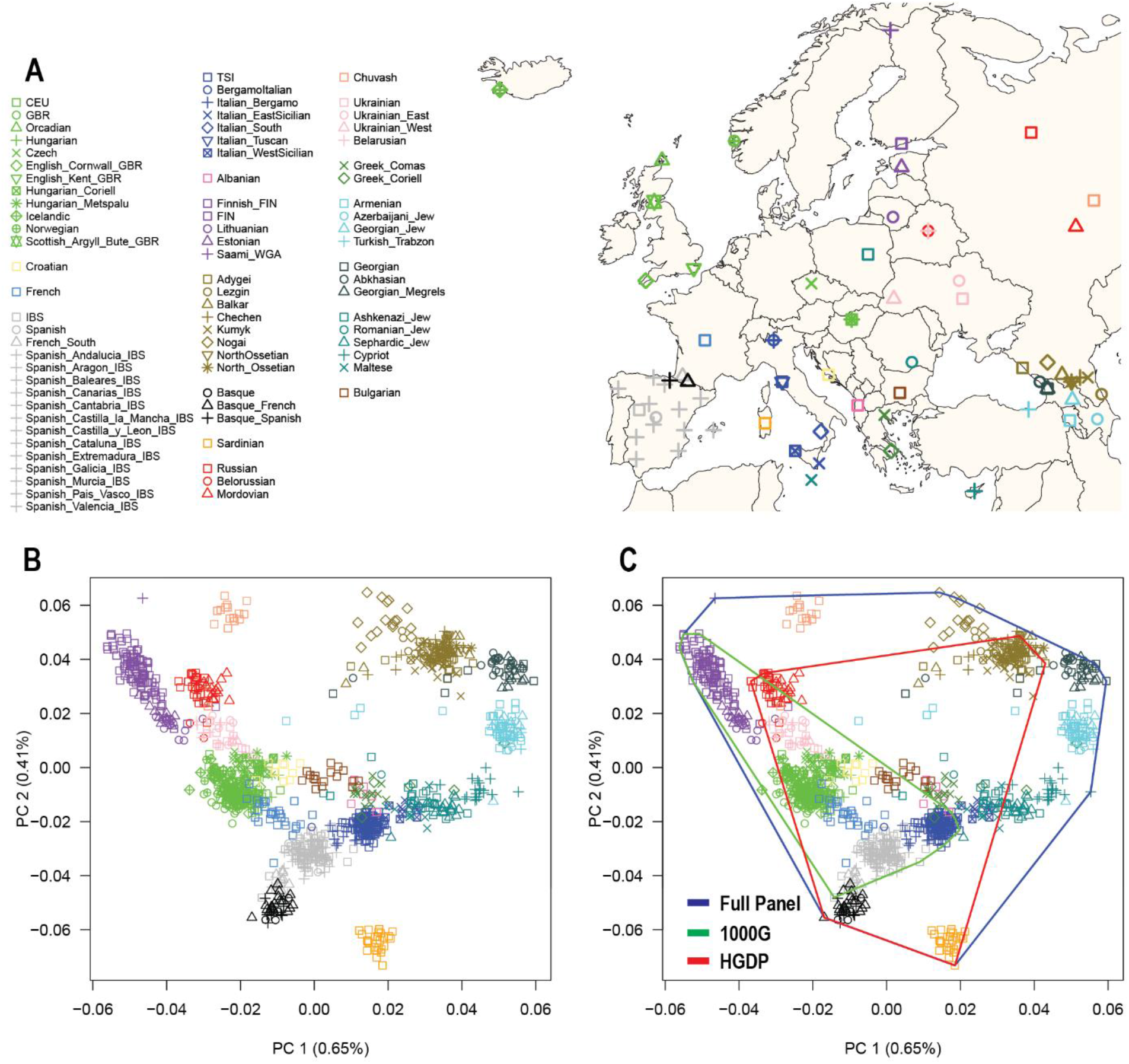
European reference panels and coverage of European genetic diversity. (A) Map of Europe showing the geographic location of samples from 79 European populations. (B) The first two principal components (PC1 and PC2) of genetic diversity and the percent variance explained. C) Coverage of genetic diversity over the first two principal components (convex hull area).

### Subcontinental stratification in individuals with European ancestry

To expand and refine our understanding of subcontinental stratification and admixture in European-ancestry populations, we integrated genome-wide genotype data from approximately 19,000 European-ancestry individuals (Fig. 2). These data included our European panel (1,216 unrelated individuals) and 17,669 European Americans from five genetic epidemiology cohorts in the US: Atherosclerosis Risk in Communities (ARIC, *n* = 9,633), Coronary Artery Risk Development in Young Adults (CARDIA, *n* = 1,675), Framingham Heart Study (FHS, *n* = 2,451), Genetic Epidemiology Network of Arteriopathy (GENOA, *n* = 1,384), and Multi-Ethnic Study of Atherosclerosis (MESA, *n* = 2,526). To assess continental-level structure, we evaluated our European-ancestry dataset with a worldwide reference panel (Fig. S2). Most Europeans formed a discrete cluster along the first two principal components, as previously observed^33,34^. Similarly, by projecting European Americans onto the worldwide reference panel, we observed that >99% of European Americans clustered with European reference individuals, with few individuals distributed along the first principal component (European-African gradient) or the second principal component (European-Asian gradient). These results suggest that the Europeans included in our panel represent a discrete cluster in relation to worldwide genetic diversity and that European Americans in genetic epidemiology cohorts in the US have small to negligible population stratification at the inter-continental scale.

**Fig. 2.**
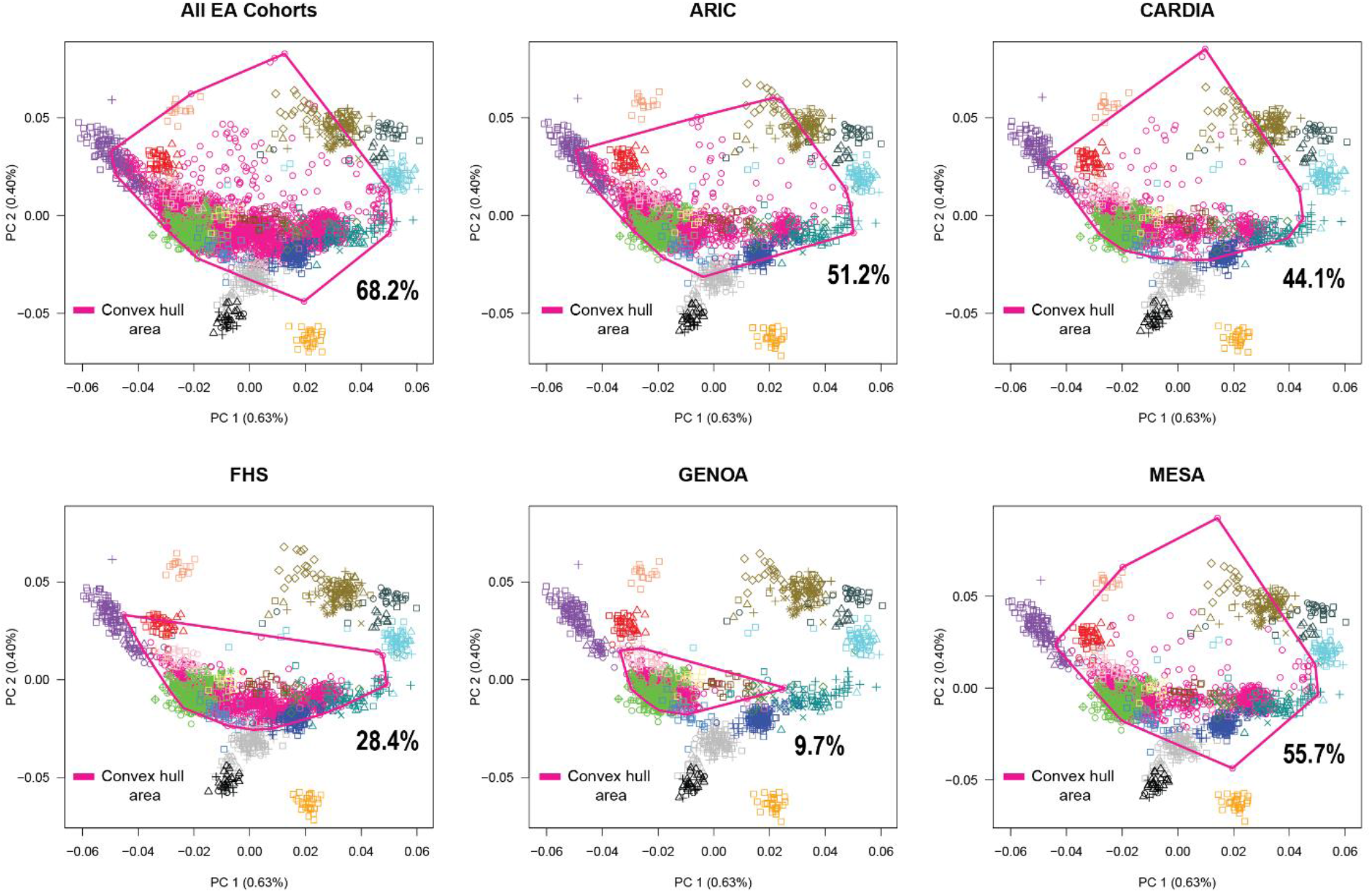
Projection analysis of European Americans onto our European reference panel. We plotted the convex hull area for all cohorts combined and for each European American cohort. The full legend as well as the geographic location of samples from 79 European populations can be found in Fig. 1. Convex hull area = Coverage of genetic diversity over the first two principal components.

Next, to evaluate European subcontinental stratification in European American cohorts, we projected individuals from each European American cohort onto our European reference panel. We calculated that European American cohorts collectively covered 68.2% of European diversity in our panel (Fig. 2), with differential coverage by cohort: 55.7% in MESA, 51.2% in ARIC, 44.1% in CARDIA, 28.4% in FHS, and 9.7% in GENOA. The ARIC, CARDIA, FHS, and MESA individuals formed at least three clusters: one with North Europeans (*e.g*., British and Scandinavian), one with Southeast Europeans (*e.g*., Ashkenazi Jew and Romanian Jew), and one overlapping Finnish individuals. GENOA individuals mostly overlapped British or Scandinavian reference individuals, with few individuals overlapping South Europeans. Most FHS samples overlapped with or were between North and South Europeans, with a large number of individuals clustering with Italian reference individuals.

### Subcontinental admixture in individuals with European ancestry

Unsupervised analysis with ADMIXTURE^17^ using our European reference panel identified the most likely number of ancestry clusters as three (Fig. 3A), suggesting that Europeans have three-way admixture among North, Southwest, and Southeast Europeans. The stacked bar plot of mixture proportions showed that the North European-associated ancestry cluster decreased along the north-to-south geographic direction (Fig. 3A). Formal correlation tests between population ancestry means and geographic coordinates revealed that latitude was significantly correlated (*p* < 2.85×10^-8^) with North European-associated ancestry cluster (Spearman’s *rho*=0.814), and longitude was correlated with Southwest- (Spearman’s *rho*=*0.859*) and Southeast-associated (Spearman’s *rho*=0.579) European ancestry clusters (Fig. 3B). We observed similar levels of genetic differentiation (*F*_ST_) between the inferred European ancestry clusters: *F*_ST_ = 0.033 between North and Southwest, *F*_ST_ = 0.032 between North and Southeast, and *F*_ST_ = 0.028 between Southwest and Southeast. To put these amounts of genetic differentiation into perspective, *F*_ST_ estimates between European ancestry clusters are comparable to *F*_ST_ between British (GBR) and either Mexican (MXL, which have ^~^50% Native American ancestry, *F*_ST_ = 0.031) or Punjabi in Pakistan (PJL, who have > 70% South Asia Ancestry, *F*_ST_ = 0.027) samples (Table S2). Additionally, *F*_ST_ estimates between European ancestry clusters are at least three-fold higher than *F*_ST_ between Mandenka from Gambia in West Africa and Luhya from Kenya from East Africa (*F*_ST_ = 0.011, Table S2). Even when comparing real-world European populations, *F*_ST_ estimates between Finnish in North Europe and Armenians or Georgians in South Europe are ^~^ two-fold higher (*F*_ST_ ^~^ 0.02) than *F*_ST_ between Mandenka and Luhya (*F*_ST_ = 0.011), *i.e*., between West and East Africans, and not as high as *F*_ST_ between inferred European ancestry clusters.

**Fig. 3.**
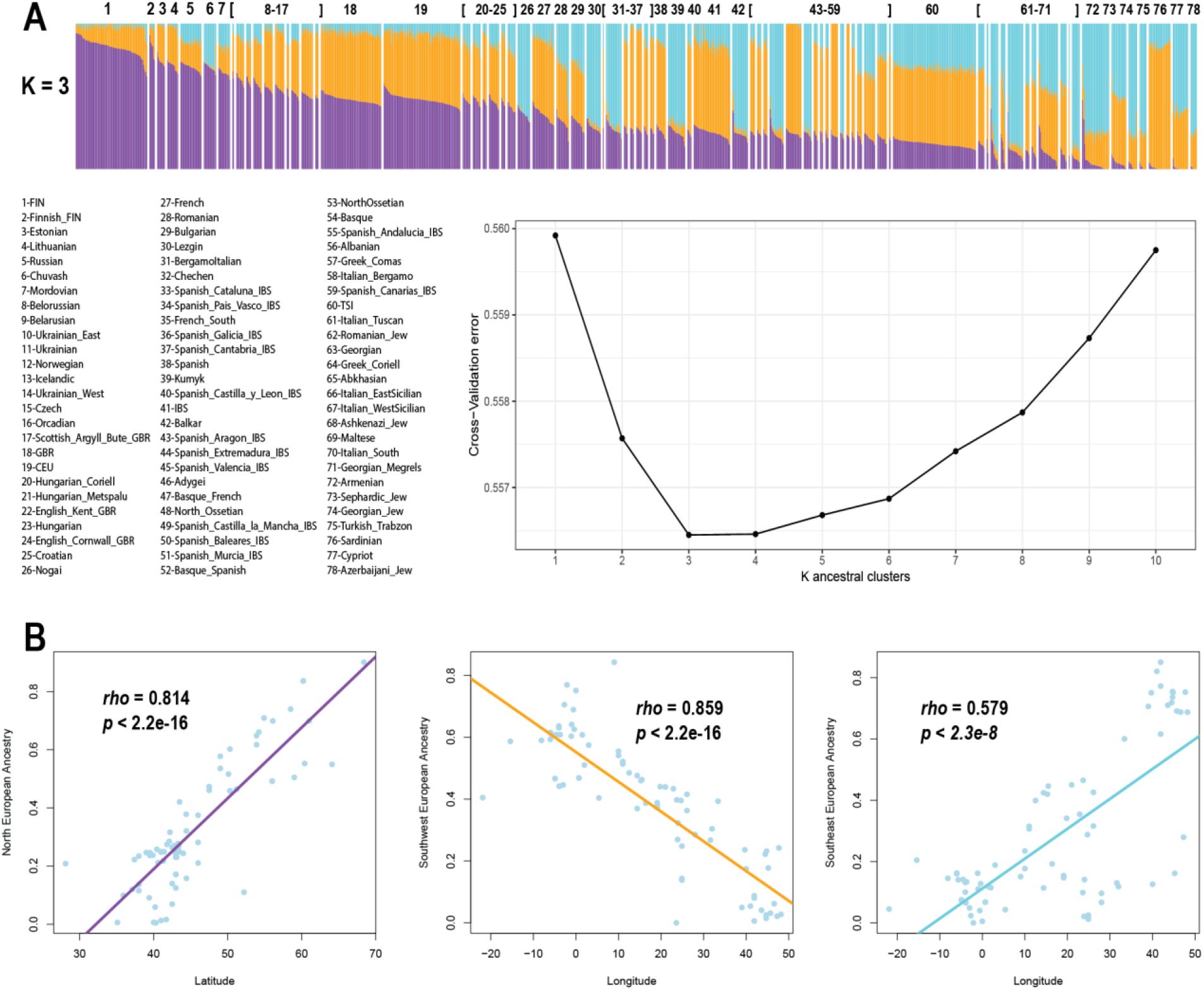
Subcontinental ancestries in Europe and correlation of ancestry with geography. (A) Bar plot showing ancestry proportions in the European populations and a cross-validation plot supporting K=3 as the most likely number of ancestry clusters. Purple, magenta, and cyan colors represent ancestry clusters associated with North, Southwest, and Southeast European populations, respectively. Individual Bar plots were sorted in descending order of the amount of North European ancestry (Purple), and populations are sorted in descending order of the average of North European ancestry. (B) Correlation plots depicting Spearman’s *rho* between ancestry proportions and geographic coordinates. Colored lines represent fitted linear regressions.

Supervised ADMIXTURE^17^ analysis of European Americans showed patterns of European ancestry clusters that differed by cohort (Fig. 4 and Table S3). GENOA had the highest mean proportion of the North European ancestry cluster (44%, SE = 3.9%) and the lowest proportion of the Southeast European ancestry cluster (7%, SE = 3%), while FHS had the lowest mean proportion of the North European ancestry cluster (29.9%, SE = 3.7%). MESA had the highest proportion of Southeast European ancestry cluster (25.4%, SE = 3.1%), followed by FHS (19.7% SE = 3%). The admixture patterns in the European American cohorts were consistent with the projection analysis (Fig. 2), *e.g*., the GENOA individuals clustered tightly with British and Scandinavian individuals on the first principal component. By testing genetic admixture using *f_3_* statistics^39^, we obtained formal evidence for admixture in the history of European Americans (Tables S4A-S4E). Also, we observed positive correlation between *F*_ST_ (a measurement of North-South European differentiation) and *F*_IT_ (a measurement of inbreeding) at SNPs throughout the genome in European American cohorts, consistent with subcontinental ancestry-related assortative mating (Table S5). Our results confirm the presence of subcontinental population structure in both Europeans and European Americans, that this structure reflects mixed ancestry in the vast majority of individuals, and that mixed ancestry reflects admixture rather than discrete subpopulations in Europe.

**Fig. 4.**
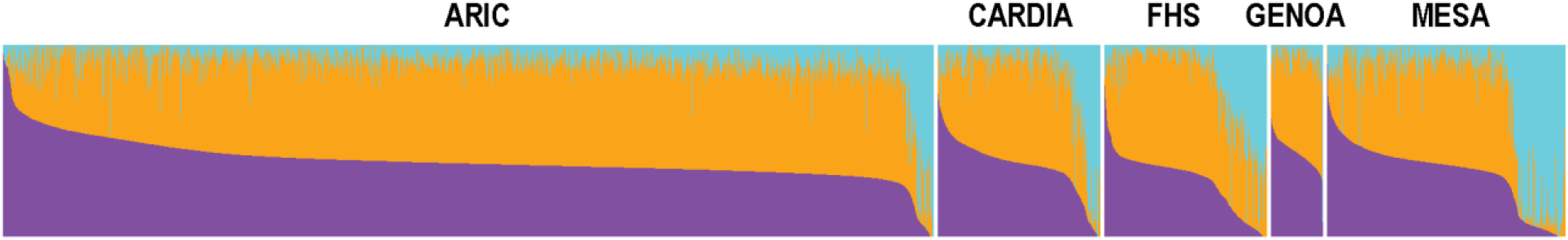
Ancestry proportions in European Americans. Bar plot representation of individual ancestry proportions inferred from supervised analysis. Purple, magenta, and cyan colors represent ancestry clusters associated with North, Southwest, and Southeast European populations, respectively. Individual Bar plots were sorted in descending order of the amount of North ancestry cluster (Purple).

### Admixture dating in European Americans

To date admixture in European Americans, we first applied a clustering approach^40^ to the first two principal components and inferred that European Americans likely cluster within three subgroups of individuals (Fig. 5A and Fig. S3). Projection analysis of European Americans onto our European reference panel revealed that European Americans were widely distributed across a north-south axis, with centroids of inferred subgroups related to North- (Subgroup N), Southwest- (Subgroup SW), and Southeast- (Subgroup SE) Europeans (Fig. 5B). The highest proportion of ancestry in Subgroup N individuals was North European ancestry (54.5%). Similarly, the highest proportions of ancestry in Subgroup SW and Subgroup SE individuals were Southwest European ancestry (53.7%) and Southeast European ancestry (71.2%), respectively. Next, we inferred admixture times for individuals within each of the three subgroups of European Americans. We observed significant admixture dating for all three subgroups, with subgroup SE yielding an admixture date ^~^10 generations more recent (42.00 generations, SE = 6.82) than admixture dates for subgroup SW (54.28 generations, SE = 10.43) and subgroup N (50.89 generations, SE = 14.26).

**Fig. 5.**
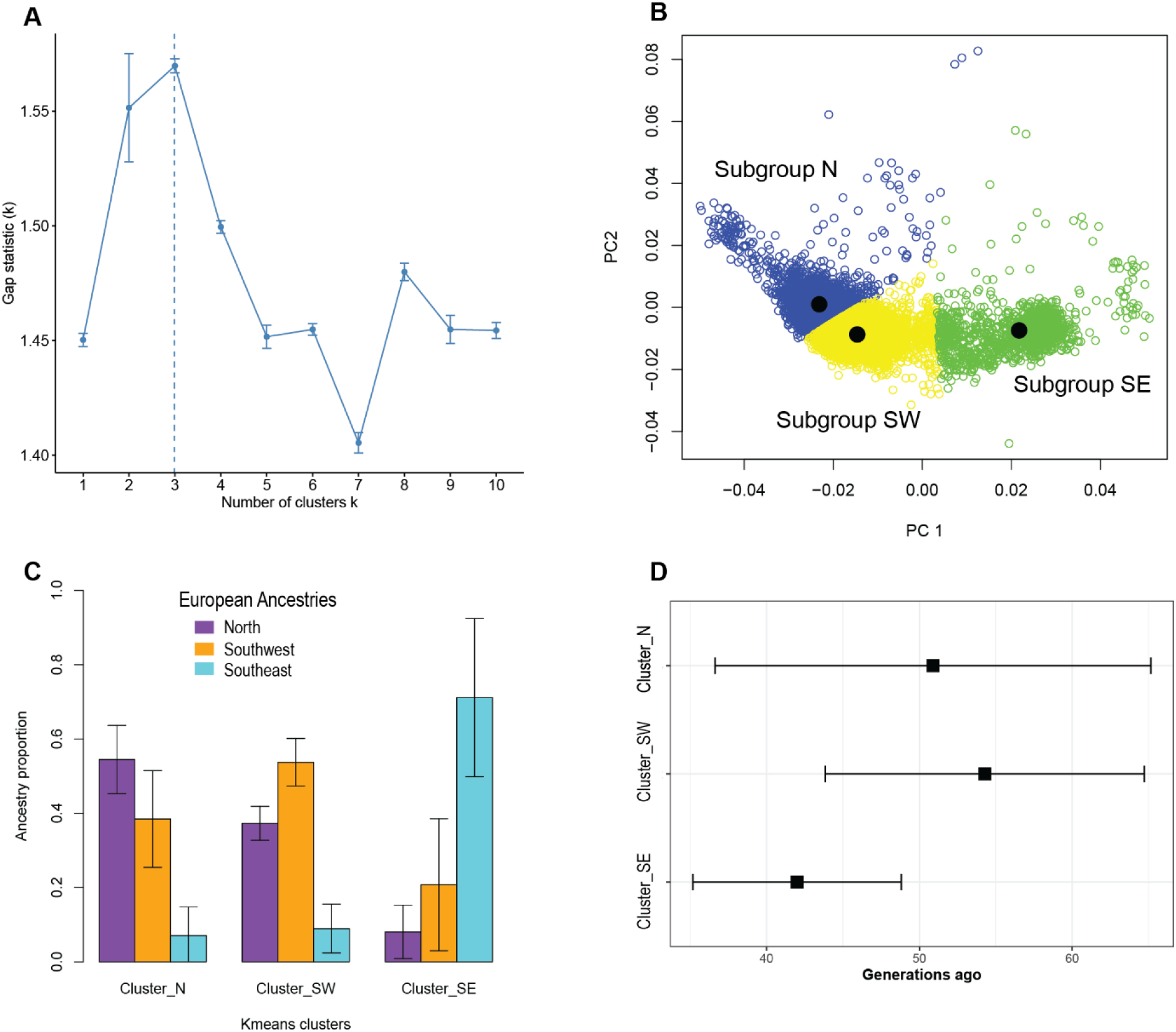
Substructure and admixture dating in European Americans. (A) The number of clusters (k) was estimated using gap statistics, based on the first two principal components (PCs) derived from the (B) projection analysis of European Americans (15,917 unrelated individuals). We estimated that European Americans are distributed across three clusters representing North (N), Southwest (SW), and Southeast (SE) Europeans. (C) Bar plot representing ancestry profiles within each estimated cluster of European Americans. D) Admixture dating across clusters of European Americans. Point estimates and standard errors of statistically significant admixture dates are shown on the horizontal axis.

### Implications of subcontinental admixture for association analysis

To understand the impact of subcontinental admixture in association studies and approaches to correct potential confounding, we investigated the classical association between *LCT* (rs4988235) and height, which has been claimed to be a false positive result due to stratification^25^. In addition, we evaluated the associations of rs4988235 with BMI and LDL, which were recently identified in large GWAS meta-analyses using primarily European-ancestry individuals (up to 500K samples)^14,23,24^. These studies either adjusted association models for genome-wide ancestry using the first 10 principal components^24^ or there was no evidence of adjustment for European population stratification^23^. Using our integrated set of European American cohorts, we replicated the previously reported associations between rs4988235 and height, LDL, and BMI when models were not adjusted for principal components, i.e., genome-wide ancestry (Fig. 6 and Table S6). Different levels of adjustment for population structure (the genetic relatedness matrix, genome-wide ancestry [PCs], and/or locus-specific subcontinental European ancestry) reduced the associations of rs4988235 with height and LDL (Fig. 6A-B and Table S6). Importantly, when models were fully adjusted for both genome-wide and locus-specific subcontinental European ancestry, the associations of rs4988235 with height and LDL were completely eliminated, indicating that the unadjusted associations were false positives. In contrast, the association between rs4988235 and BMI remained weakly significant after adjustment for both genome-wide and locus-specific ancestry (Fig. 6C and Table S6).

**Fig. 6.**
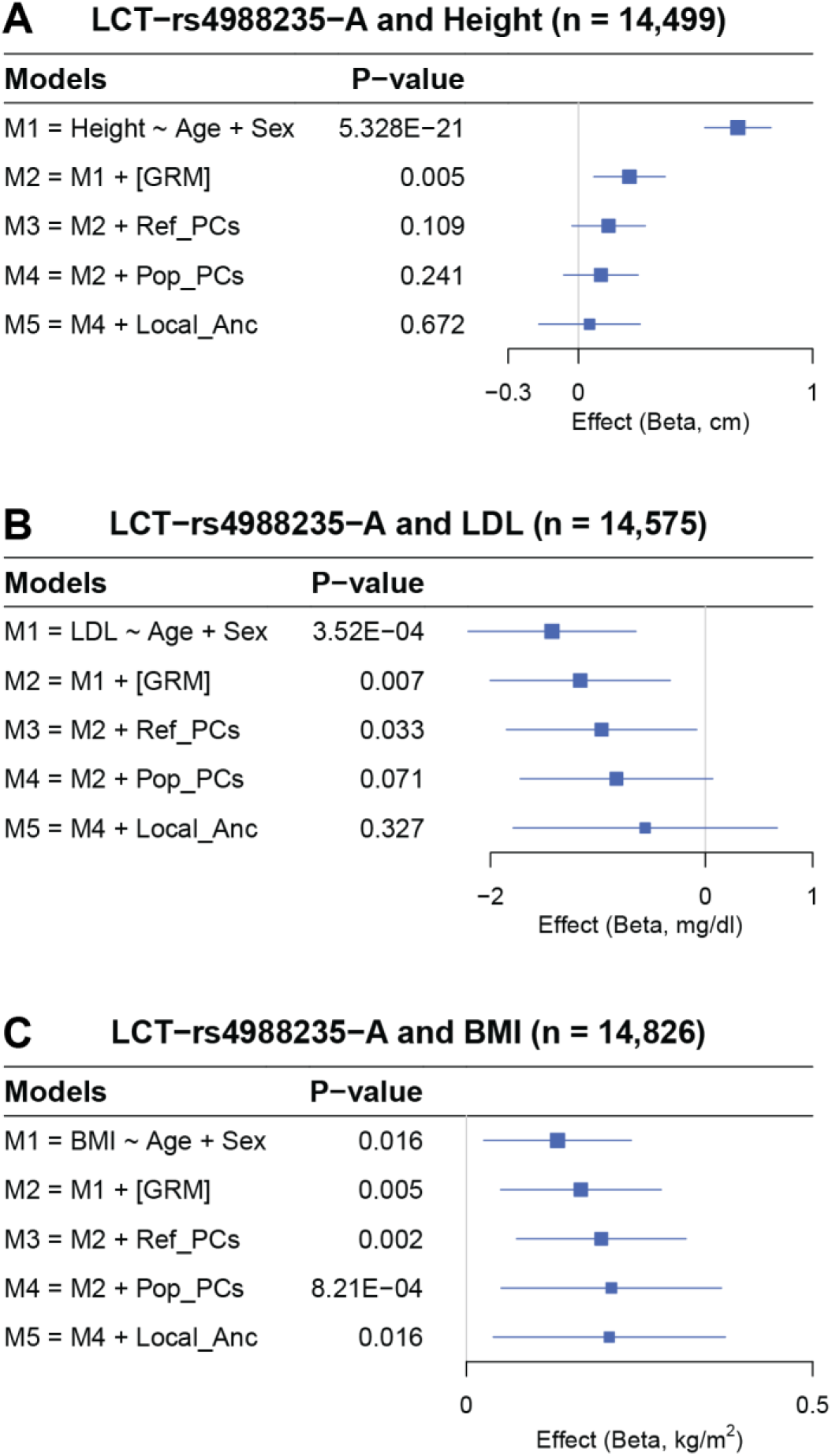
Forest plots showing the association between rs4988235 and height, LDL, and BMI, accounting for different levels of control of population stratification. Forest plots show β values (95% confidence intervals) and *p*-values from linear mixed models. GRM = genetic relatedness matrix; Ref_PCs = PCs derived from a projection of individuals onto an ancestral reference panel; Pop_PCs = PCs derived from within-population unsupervised PCA analysis.

We also performed cohort-specific association analysis between rs4988235 and height, BMI, and LDL (Tables S7-S9). When models were not adjusted for population stratification, the association between rs4988235 and height was significant in ARIC, CARDIA, FHS, and MESA but not in GENOA (Table S7). The lack of association in GENOA might be explained by a small amount of ancestral heterogeneity and/or by small sample size. After adjustment for genome-wide ancestry, we observed association between rs4988235 and height in CARDIA but not in the other four cohorts. After adjustment for genome-wide and locus-specific ancestry, we observed no association between rs4988235 and height in all five European American cohorts (Table S7). Similarly for LDL, we observed some cohort-specific associations when models were not fully corrected, and that full adjustment reduced or eliminated significance in all cohorts (Tables S8 and S9). These results imply that full ancestry adjustment (genome-wide and locus-specific subcontinental ancestry) may facilitate correction for residual stratification and avoidance of false positives in single studies.

It is common practice in genetic association studies to account for genome-wide ancestry using principal components derived from study-specific unsupervised analysis (population-specific PCA). Here, we tested the approach of deriving principal components from projection of target individuals onto an external reference panel (projection or supervised PCA). To evaluate the similarity between these two approaches using our European American data, we performed Mantel’s correlation test between individuals’ genetic distances computed from the top twenty principal components obtained from the unsupervised and projection approaches. We observed moderate correlation in four studies (Mantel’s *rho* from 0.46 to 0.53, *p* < 0.001), with GENOA not showing a significant correlation (Table S5). Differences between these two PCA approaches may have led to differences in how well confounding was controlled. During testing of the association between rs4988235 and height, we observed systematically better model fits (*Δ*AIC up to 12.45)^41^ across cohorts when models were adjusted for projection-derived principal components compared to study-specific principal components (Table S7). For the integrated data set, projection-derived and study-specific principal components provided similar model fits (Table S6).

## DISCUSSION

The existence of subcontinental-level ancestries has been documented within Africa and Asia^4,42–44^, yet the presence of European subcontinental ancestries within Europe is not well appreciated. We compiled genome-wide genotype and sequence data from geographically diverse Europeans and European Americans to investigate subcontinental-level ancestries and admixture in European-ancestry individuals. We also explore the consequences of different strategies for addressing ancestry in genetic epidemiology studies. Our study has four major results, described below.

First, we created a new reference panel of European genetic diversity by combining five genome-wide data sets^33–37^. We showed that panels based on the 1000 Genomes Project and the Human Genome Diversity Project, separate or combined, provided incomplete coverage of genetic diversity among Europeans or the European component of European Americans compared to our new reference panel. To facilitate genome-wide ancestry estimates, we provide as a research resource a reference SNP matrix of subcontinental ancestry-specific allele frequencies (https://github.com/mateushg1/CRGGH/). This resource allows for estimation of subcontinental ancestry proportions by projection analysis based on publicly available, aggregated, and non-identifiable data. The end-user does not need to access, clean, integrate, or analyze individual-level reference data.

Second, our admixture analyses yielded formal evidence that European-ancestry individuals are admixed at the subcontinental level. Using multiple approaches to infer admixture, we showed that European-ancestry individuals are three-way admixed with wide variation in ancestry proportions. The demonstration that European Americans are ancestrally heterogeneous has implications for calibrating locus-specific ancestry analysis^19^ with respect to the number of generations since admixture began. Admixture dates estimated for European Americans corresponded to the large-scale Migration Period in Europe (300-800 AD)^45^, and were consistent with gene flow after the end of Roman Empire described in ancient DNA studies of the Viking Age^11^ and Anglo-Saxon migrations^12^. Moreover, our results support the occurrence of subcontinental ancestry-related assortative mating as a social factor that shaped the genetic structure of European Americans in the US^46^.

Third, studies of European-ancestry individuals have reported that genetic variants, principally rs4988235, in the lactase gene (*LCT*) are associated with height, BMI, and LDL^23,24,47^. However, the association between rs4988235 and height has been suggested to be spurious due to uncorrected genome-wide ancestry^25^. Adjustment for genome-wide ancestry may not be sufficient to avoid false positive results and can mask true associations if ancestry is associated with the outcome^48^. Consistent with known potential confounding effects of ancestry^3,49^, we demonstrated that the lack of adjustment for both genome-wide and locus-specific ancestry can produce false positives in association studies using European-ancestry individuals. By adjusting our models for locus-specific ancestry in addition to genome-wide ancestry, associations of rs4988235 with height and LDL were eliminated. In contrast, the association between rs4988235 and BMI remained after correcting for both genome-wide and locus-specific ancestry, suggesting an effect on weight but not on height. These results suggest that residual confounding by subcontinental European ancestry can produce spurious associations in genetic association studies, with consequences for polygenic adaptation^15^, polygenic risk scores^16^, and fine-mapping of genetic associations. Importantly, our results indicate that adjustment for subcontinental European ancestry at both genome-wide and locus-specific levels should be considered in genetic association studies using European-ancestry individuals, including large biobanks such as the UK Biobank^30^ in Europe and the All of Us Research Program^31^ and the Million Veteran Program^32^ in the United States.

Fourth, we observed better model fit with adjustment for principal components derived from supervised analysis based on a common reference panel rather than for principal components derived from study-specific unsupervised analyses. However, the performance of unsupervised analysis approached the performance of supervised analysis as the genetic diversity covered by the sample data approached the genetic diversity covered by the external reference panel. European genetic diversity in our full panel covered by European American cohorts ranged from 9.7% to 55.7% whereas coverage reached 68.2% when all cohorts were combined. This result indicates that GWAS meta-analyses in which individual-level data cannot be or are not shared across studies should rely on supervised analysis given a common reference. This recommendation does not depend on sample size, as even data sets on the scale of large biobanks do not necessarily cover a large proportion of ancestral diversity.

In conclusion, we demonstrated that European-ancestry individuals are admixed at the subcontinental level. Subcontinental admixture in Europeans and European Americans, if not properly accounted for, can produce false positive associations in genetic epidemiology studies due to incomplete correction for confounding by ancestry. Our study highlights the need for full control, at both genome-wide and locus-specific ancestry levels, for confounding in Europeans and European Americans. Potential consequences of residual confounding by subcontinental ancestry include the misestimation of polygenic adaptation and poor performance of genetic or polygenic risk scores.

## METHODS

### Samples

We compiled genome-wide data from five different studies: the 1000 Genomes Project^33^, the Human Genome Diversity Project (HGDP)^34^, the Human Origins dataset^35^, a study of the Caucasus Mountains^36^, and a study of the Jewish Diaspora^37^ (Fig. 1A and Table S1). Using these data, we created a data set that included 4,796 individuals (worldwide reference panel), from which we extracted 1,216 individuals from 79 European populations (European reference panel). We analyzed genome-wide array and phenotypic data from 17,684 European Americans from five genetic epidemiology cohorts, for which access was granted through dbGaP^50^: ARIC (phs000090.v1.p1), CARDIA (phs000285.v3.p2), FHS (phs000007.v32.p13), GENOA (phs000379.v1.p1), and MESA (phs000209.v13.p3).

### Data curation

To reduce batch effects due to the integration of array-based genotype data and whole genome sequence data, we performed quality control analysis within and between datasets using PLINK 1.9, filtering by minor allele frequency (*--maf 0.01*), per genotype missingness (*--geno 0.05*), per individual missingness (*--mind 0.05*), and deviation from Hardy Weinberg equilibrium (*--hwe 1×10^-6^*). We also pruned strand-ambiguous SNPs and SNPs in high linkage disequilibrium (*--indep-pairwise 50 10 0.8*).

### Population structure and relatedness

We used PLINK 1.9 to estimate the probability that individuals *i* and *j* share 0, 1, or 2 alleles identical by descent (IBD) (δ^0^_*ij*_, δ^1^_*ij*_, and δ^2^_*ij*_, respectively)^51^. Based on these IBD probabilities, we calculated the pairwise kinship coefficient (*Φ_ij_*) as a function of IBD-sharing, *Φ_ij_* = *1/2*δ^2^_*ij*_ + *1/4*δ^1^_*ij*_. We modeled the genetic relationships among individuals as networks^52^, in which pairs of individuals were linked if they had a *Φij* threshold ≥ 0.0884 (*i.e*., first- and second-degree relatives^53^). Then, we excluded related individuals using the maximum clique graph approach to minimize sample loss^52^. We performed unsupervised principal components analysis^13^ and unsupervised ADMIXTURE analysis^17^ on the European reference data. We performed unsupervised and supervised PCA and ADMIXTURE analyses using the reference data combined with the European American data. For supervised analysis in ADMIXTURE, we used as the ancestral references the European individuals with ≥90% of one of three ancestries based on unsupervised ADMIXTURE analysis. To evaluate the coverage of European diversity, we used the first two principal components to calculate convex hull areas^38^. We calculated *f*_3_ statistics as implemented in ADMIXTOOLS^39^ to formally test admixture. We tested all possible combinations of two European sources and a target European American cohort, following the form *f*_3_(EUR_POP_X, EUR_POP_Y; EA_Cohort). All *f*_3_ statistics with *z* ≤ −3 were considered significant evidence of admixture.

### Admixture dating

We first combined all European American cohorts and performed supervised PCA by projecting the European Americans onto the European reference panel. We then used gap and elbow statistics^40^ to calculate the most likely number of clusters. To estimate the origin dates of admixture events, we calculated weighted LD decay statistics using MALDER^54^ within each cluster of European Americans. Given that background LD can have a confounding effect on the weighted LD curves, we used as reference populations North European (Lithuanian and Estonian) and South European (Cyprus, Azerbaijani Jew, and Georgian Jew) populations that did not show high LD correlation with the tested target populations.

### Phasing and imputation

To generate valid VCF files before phasing, imputation, and association tests, we checked and corrected for monomorphic sites, consistency of reference alleles with the reference genome, variants with invalid genotypes, and non-SNP sites using the checkVCF.py Python script (https://github.com/zhanxw/checkVCF). We phased and imputed the genotype data using EAGLE2.4^55^ and Minimac^56^, respectively, using the TOPMed panel available through the TOPMed imputation server^57^. After imputation, we retained SNPs with minor allele frequency ≥0.01 and with either high imputation quality (info ≥ 0.95) or empirically determined genotype data.

### Locus-specific ancestry analysis

Given that rs4988235 is highly differentiated between North and South European populations^28^ and varies following a north-to-south gradient^25^, we inferred two-way locus-specific ancestry using RFMix (version 1.5.4)^19^. For ancestral references, we selected individuals with ≥90% North or South European ancestry as estimated in the unsupervised ADMIXTURE analysis. We performed inference in the PopPhased mode to correct possible phase errors. We set the number of generations since the admixture event (argument - G) at 50, the number of expectation maximization (EM) iterations (argument -e) at 2, and the window size (argument -w) at 0.2 cM. All other arguments were set at default values.

### Association analysis

We performed association analysis using linear mixed models implemented in GENESIS^58^. Our analyses were focused on unrelated European Americans, with relatedness determined by the maximum clique graph approach^52^. Models were adjusted for the genetic relationship matrix as a random effect to account for variance components and the four first principal components (significantly associated with the outcome) and/or locus-specific ancestry as fixed effects. genome-wide ancestry was accounted for using principal components derived from one of two approaches: study-specific unsupervised analysis or supervised (projection) analysis of individuals onto an external reference panel. To account for the uncertainty of locus-specific ancestry estimates, models were adjusted for locus-specific ancestry dosages calculated from the posterior probabilities of locus-specific ancestry. Similarly, we used genotype dosages to account for imputation uncertainty.

## ETHICS STATEMENT

All dbGaP studies (dbGaP Study Accession described in the Methods section) obtained ethical approval from the relevant institutions and written informed consent from each participant prior to participation. We obtained approval for controlled access (protocol number: 12-HG-N185) of each of the dbGaP studies.

## ACKNOWLEDGMENTS

We are thankful to the participants in the ARIC, CARDIA, FHS, GENOA, and MESA, their families, and their physicians. The High Performing Computation (HPC) group at the National Institutes of Health. We thank Dr. Thiago Peixoto Leal to provide support to local ancestry pipelines on GitHub, and scientific discussions. The views expressed in this manuscript are those of the authors and do not necessarily represent the views of the NIH.

## AUTHOR CONTRIBUTIONS

The project was conceived by M.H.G., D.S., C.N.R., and A.A.A. M.H.G. and D.S. assembled datasets. M.H.G. and D.S. analyzed genetic data. M.H.G., D.S., A.R.B., E.T.S., C.D.B., A.A.A., and C.N.R. contributed to data interpretation. M.H.G., D.S., A.A.A., and C.N.R wrote the manuscript. All authors read the manuscripts and contributed with suggestions.

## Notes

### Competing Interest Statement

The authors have declared no competing interest.

### Summary of Updates

The author's List was revised with the middle names included

