## Supplemental Figures for "Unappreciated Subcontinental Admixture in Europeans and European Americans: Implications for Genetic Epidemiology Studies"

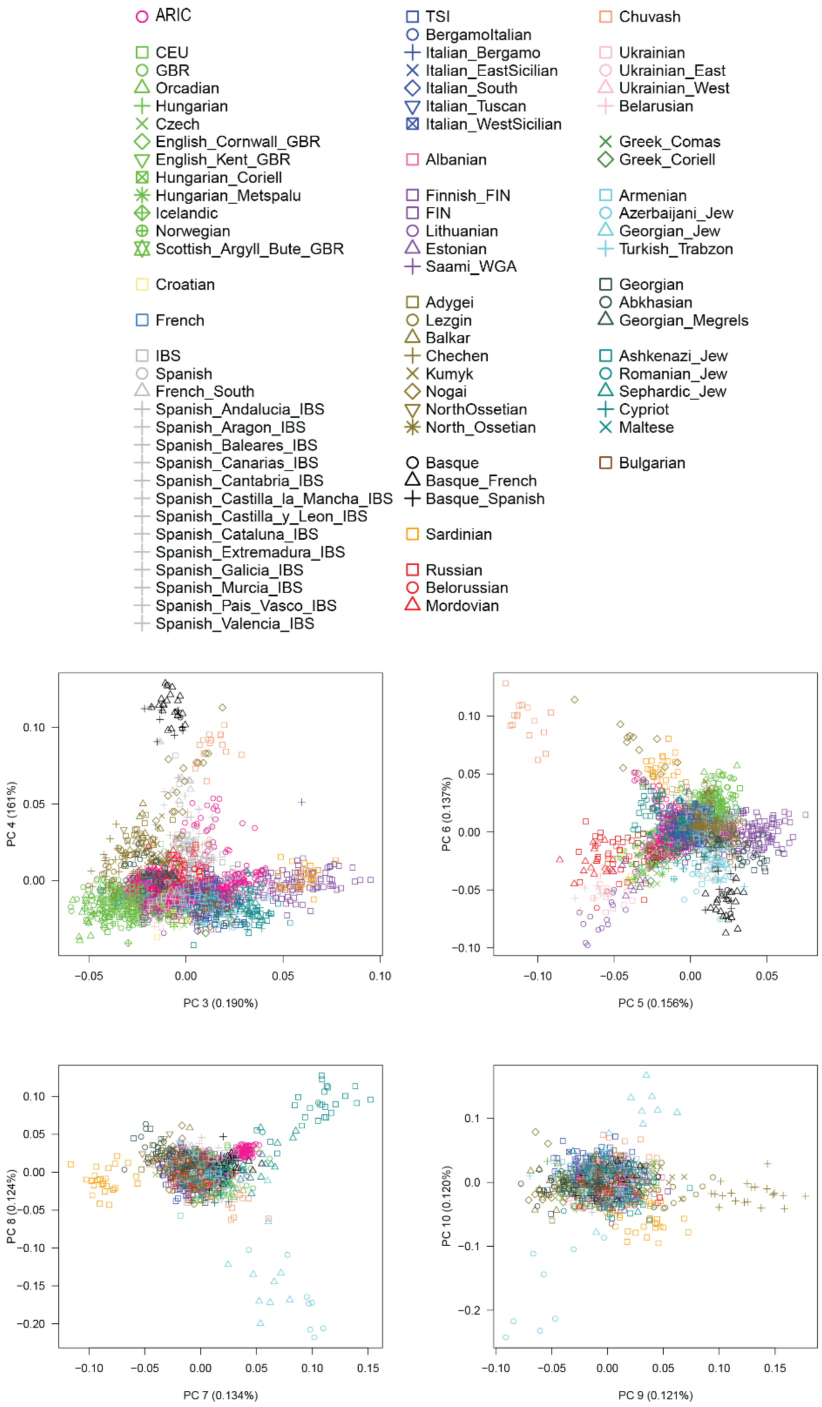


**Fig. S1. Principal component analysis (principal components 3 through 10) of the European reference panel.** European Americans (ARIC cohort) were projected onto the European reference panel.

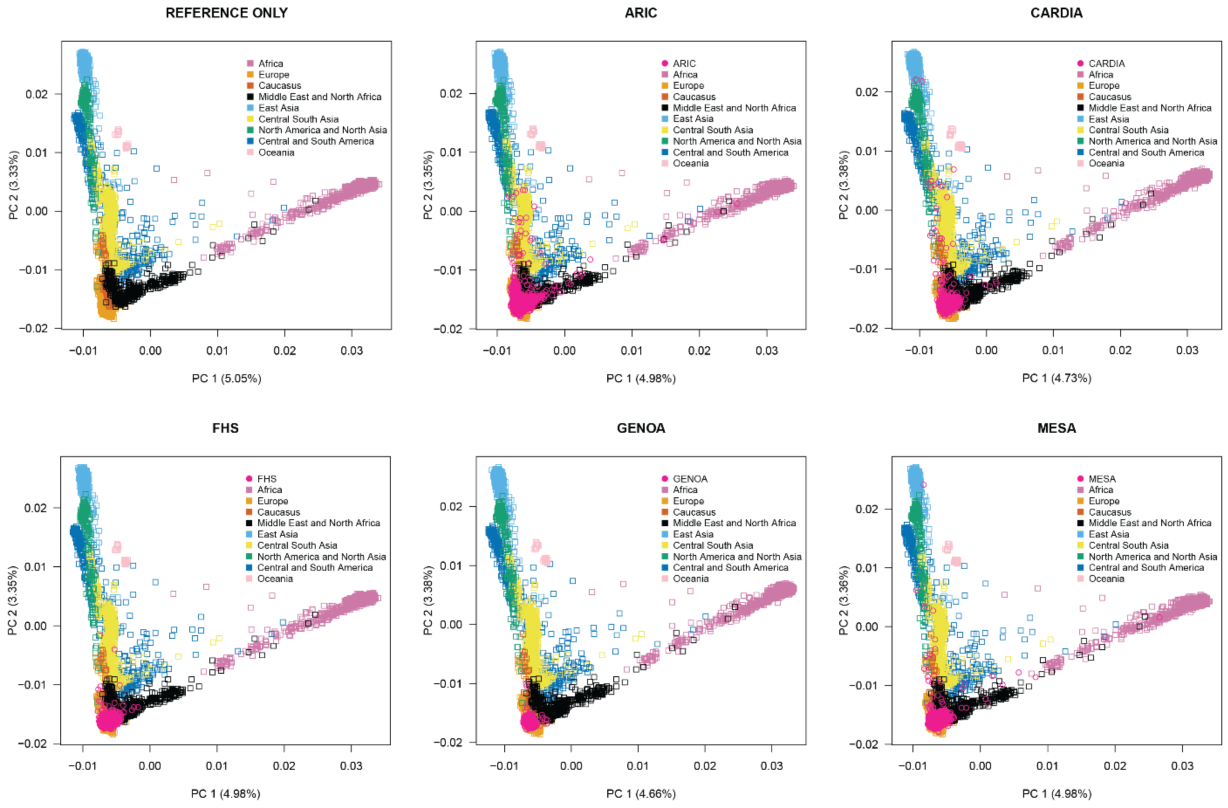


**Fig. S2. Principal component analysis of the worldwide reference panel.** Each European American cohort is projected onto the worldwide reference panel.

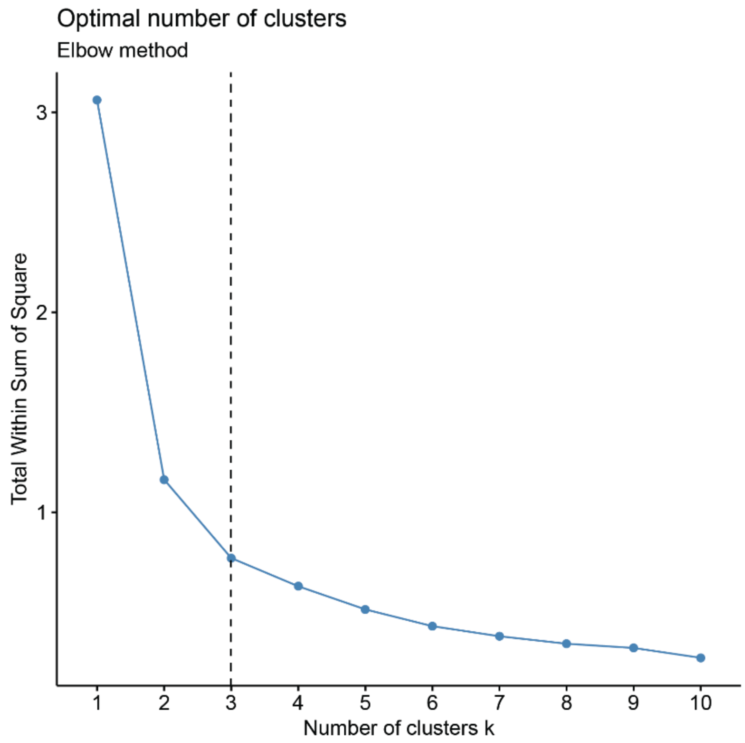


**Fig. S3. Estimated number of clusters (k) using the elbow method.** The number of clusters of European American individuals was estimated from the first two principal components derived from the projection of all European Americans onto the European reference panel.
